# Beyond Alternans: Detection of Higher-Order Periodicity in Ex-Vivo Human Ventricles Before Induction of Ventricular Fibrillation

**DOI:** 10.1101/2023.05.01.539003

**Authors:** Shahriar Iravanian, Ilija Uzelac, Anand D Shah, Mikael J Toye, Michael S. Lloyd, Michael A. Burke, Mani A Daneshmand, Tamer S Attia, J David Vega, Faisal M. Merchant, Elizabeth M Cherry, Neal K. Bhatia, Flavio H. Fenton

## Abstract

**Background:** Repolarization alternans, defined as period-2 oscillation in the repolarization phase of the action potentials, is one of the cornerstones of cardiac electrophysiology as it provides a mechanistic link between cellular dynamics and ventricular fibrillation (VF). Theoretically, higher-order periodicities (e.g., period-4, period-8,…) are expected but have very limited experimental evidence.

**Methods:** We studied explanted human hearts, obtained from the recipients of heart transplantation at the time of surgery, using optical mapping technique with transmembrane voltage-sensitive fluorescent dyes. The hearts were stimulated at an increasing rate until VF was induced. The signals recorded from the right ventricle endocardial surface just before the induction of VF and in the presence of 1:1 conduction were processed using the Principal Component Analysis and a combinatorial algorithm to detect and quantify higher-order dynamics.

**Results:** A prominent and statistically significant 1:4 peak (corresponding to period-4 dynamics) was seen in three of the six studied hearts. Local analysis revealed the spatiotemporal distribution of higher-order periods. Period-4 was localized to temporally stable islands. Higher-order oscillations (period-5, 6, and 8) were transient and primarily occurred in arcs parallel to the activation isochrones.

**Discussion:** We present evidence of higher-order periodicities and the co-existence of such regions with stable non-chaotic areas in ex-vivo human hearts before VF induction. This result is consistent with the period-doubling route to chaos as a possible mechanism of VF initiation, which complements the concordant to discordant alternans mechanism. The presence of higher-order regions may act as niduses of instability that can degenerate into chaotic fibrillation.

## Introduction

Malignant ventricular arrhythmias, including polymorphic Ventricular Tachycardia (VT) and Ventricular Fibrillation (VF), are the proximate cause of death in many patients. Therefore, understanding the dynamics of VF and its initiation is of utmost theoretical and practical importance.

VF is the chaotic consequence of multiple drifting and interacting reentrant waves.^1–3^ Here, we use *chaos* in the nonlinear dynamics sense, i.e., “*an aperiodic long-term behavior in a deterministic system that exhibits sensitive dependence on initial conditions*”.^4^ The route from organized cardiac activity to the chaotic VF has yet to be fully explained. One leading hypothesis is based on the concept of dynamically induced dispersion of repolarization alternans.

Action Potential Duration (APD) alternans is defined as the beat-to-beat (period-2) oscillation in the APD and is the simplest quantifier of repolarization dynamics.^5^ Most cardiac tissues exhibit APD alternans when stimulated at a sufficiently fast rate. It is relatively easy to initiate and detect APD alternans in various experimental models. APD alternans can even be detected in clinical settings as the microvolt T-wave alternans (TWA), recorded using specialized ECG systems.^6^ Historically, TWA has been used for the risk stratification of malignant ventricular arrhythmias.^7^ Additionally, the possibility of feedback control of alternans using timely cardiac stimulation has been extensively studied.^8–10^ However, the practical applications of APD alternans have fallen from favor and are rarely discussed nowadays. Among many reasons, low specificity (low positive predictive value) of alternans detection in predicting malignant ventricular arrhythmias has been a significant obstacle.^11^ Despite this, APD alternans has maintained a central place in the study of cardiac dynamics and the mechanisms of VF.

Multiple mechanisms have been considered for generating APD alternans. These mechanisms can broadly be categorized as voltage-driven or calcium-driven.^5,12^ The main voltage-driven mechanism is based on the restitution hypothesis that posits that the APD is a function of the previous diastolic (resting) interval. The restitution curve, the relationship between the APD and the diastolic interval, is usually monotonically increasing and can be shown to induce alternans at fast rates, where the slope of the curve is above 1.^13^ Subsequent experiments detected alternans within regions of the parameter space not predicted by the restitution hypothesis, as in reality, the APD restitution is a multidimensional function depending on more than the previous diastolic interval. The nonlinear excitation-contraction coupling (the calcium-driven mechanism) is the second mechanism for alternans. During normal cardiac excitation, calcium entry through the sarcolemma channels triggers additional calcium release from the intracellular stores, which should cycle back into the sarcoplasmic reticulum before the subsequent beat. When the heart is paced fast or subjected to pathological conditions (e.g., ischemia, heart failure), the calcium recycling mechanism may lag compared to the transmembrane potential and is prone to oscillation and complex and potentially chaotic behavior.^14,15^

The primary reason to study alternans is its mechanistic link to the generation of unstable substrates that are precursors to VF. The established hypothesis is based on the concordant to discordant alternans mechanism of VF. When a sufficiently large segment of the heart is stimulated at a slow pacing rate, action potential waves will propagate with no discernible beat to beat variation in their duration. As the stimulation cycle length decreases, the dynamics eventually undergoes a bifurcation such that period-2 oscillation in APD starts. Initially, the APD alternans has low amplitude, and the entirety of the ventricles alternates in phase (all regions have long APDs on the same beat and short APDs on the next beat). This condition is called *concordant alternans* and is not directly proarrhythmic. As the ventricles are stimulated at faster rates, subtle variations in conduction velocity interact with the APD dynamics and cause *discordant alternans* to emerge when different regions alternate out of phase.^16^ Alternans can produce large dispersions in the APD leading to functional conduction block, specially following the onset of alternans in the action potential amplitude,^17^ or can interact with an extra stimulus within a vulnerable period (say, a premature beat), both of which can lead to the initiation of VF.^16,18,19^

A main characteristic of VF initiation through the discordant alternans route is the clear demarcation between the absence of chaos at the discordant alternans stage and the presence of chaos as soon as VF initiates. In this paper, our goal is to study an alternative mechanism of VF initiation based on the classic period-doubling route to chaos.^20^ Under this route, the classic APD alternans is a period-2 rhythm, resulting from the first period doubling. However, period doubling does not stop at period-2. Specially, excitation-contraction coupling can generate higher-order periodicities. We anticipate that as the cycle length becomes shorter, higher order periods (1:4, 1:8, 1:16,…) appear until the system transitions to chaos. The resulting chaotic activities are initially regional and contained (i.e., chaos coexists with regular activity for a period of time) but may eventually spill over and capture the whole heart, resulting in VF.

The intermediate stages in the period-doubling cascade in cardiac tissue have been experimentally elusive and are reported in only a few animal models. Savino et al. reported higher-order oscillations in bullfrog ventricles.^21^ Gilmour et al. showed period-16 and chaotic behavior in canine cardiac Purkinje fibers.^22^ Gizzi et al. demonstrated period-4 and period-8 in canine right ventricular preparations.^23^ The usual explanation for the difficulty of observing higher-order dynamics is that the width of the regions at which the next period doubling develops decreases exponentially according to the Feigenbaum constant. In this paper, we report on the detection of low-amplitude stable period-4 and higher-order oscillations in human hearts.

## Methods

### Heart Harvesting

The study protocol was approved by the Emory University and Georgia Institute of Technology Institutional Review Boards (IRB). We obtained hearts from the recipients of orthotopic heart transplantation at the Emory University Hospital. The patients consented to the research protocol before the surgery. At the time of surgery, each patient was fully heparinized and placed on a cardiopulmonary bypass machine after circulatory arrest was induced with the infusion of cold cardioplegia solution. Then, the pericardium was opened, and the recipient’s heart was removed using the bicaval technique.

Within 5 minutes of heart harvesting, the explanted heart was perfused from the arteries with cold cardioplegia solution for 5 minutes and transported to the optical mapping lab. Once in the lab, the left main and right coronary arteries were cannulated, and the heart was perfused with warm (37C) oxygenated Tyrode’s solution until the return of the spontaneous contractions. The heart was placed in an imaging chamber and perfused for at least half an hour to recover from the cardioplegia. Before optical mapping, the cardiac motion was suppressed with the help of the myosin ATPase inhibitor (-)-Blebbistatin, at a concentration of 1.8 uM.^24^

### Optical Mapping

One of the main tools to study complex arrhythmias is optical mapping.^25^ Staining arterially-perfused explanted whole heart or a segment of it with voltage- and calcium-sensitive fluorescent dyes allows for non-contact mapping of the transmembrane potential and intracellular calcium concentration with high spatial (sub-millimeter) and temporal (milliseconds) resolutions.

In this study, after the heart was prepared as above (arterially cannulated, perfused, and immobilized), it was stained with 1 mg of near-infrared voltage-sensitive dye JPW-6003 (also known as di-4-ANBDQPQ) dissolved in ethanol.^26,27^ Some hearts were also stained with a calcium-sensitive dye. The first voltage and calcium measurements were obtained from the epicardial surface as long as there was an acceptable imaging window (depending on the amount of fat across the epicardial surface). Afterward, the right ventricle free wall was removed, and its endocardial surface was imaged. We report on the results of transmembrane potential mapping from the endocardial surface of the right ventricles.

The tissue was excited using a deep red LED coupled with a 660/20 nm bandpass filter, and the emitted fluorescence was directed through a long pass 700 nm filter into an Electron-Multiplying-Charged-Coupled-Camera (EMCCD) camera. Image acquisition was performed at 500 Hz and with a resolution of 128 × 128 pixels.

### Study Protocol

The study protocol was organized as multiple restitution runs. In each run, the right ventricles were paced at progressively faster rates, starting at a pacing cycle length of ∼2000 ms and down to either the emergence of 2:1 block or induction of a reentrant arrhythmia (VT or VF). Optical signals were recorded from the endocardial surface in 20-40 seconds segments. To detect the 1:4 peak, we compared the signal recorded just before VT/VF induction (while still global 1:1 capture and conduction were present) to a control recording at 500 ms.

### Signal/Image Processing

The transmembrane potential data for each pixel is low-passed filtered by a cutoff of 50 Hz, and the output is normalized in the 0 to 1 range. Traditionally, the next step in most optical mapping studies is spatial filtering by convoluting the signal at each point in time with a suitable spatial kernel (usually a boxcar or Gaussian kernel). In this paper, we use a variational method to perform spatial filtering to better control and improve the definition of wavefronts (Supplement A).

Period-4 and higher-order dynamics are not distributed uniformly over the recording area.^23^ Instead, they are localized to a few regions. In response, we have developed two complementary processing pathways, one *global* to detect the presence of the low-amplitude period-4 and higher signals from the whole imaging area and the other *local* to localize period-4 and higher at a pixel level. The local algorithm has high sensitivity and is paired with the high-specificity global algorithm to reduce spurious detection of higher periodicity.

The key to global analysis is to remove the effects of wave propagation to distill the dynamics to a few aggregate channels focused on the repolarization phase. We shift the signals to align the upstrokes (Figure 1, panel A is unshifted signals and B is frame shifted). Because our signals were recorded while pacing the heart at a stable rate, frame shifting is possible.

**1.**
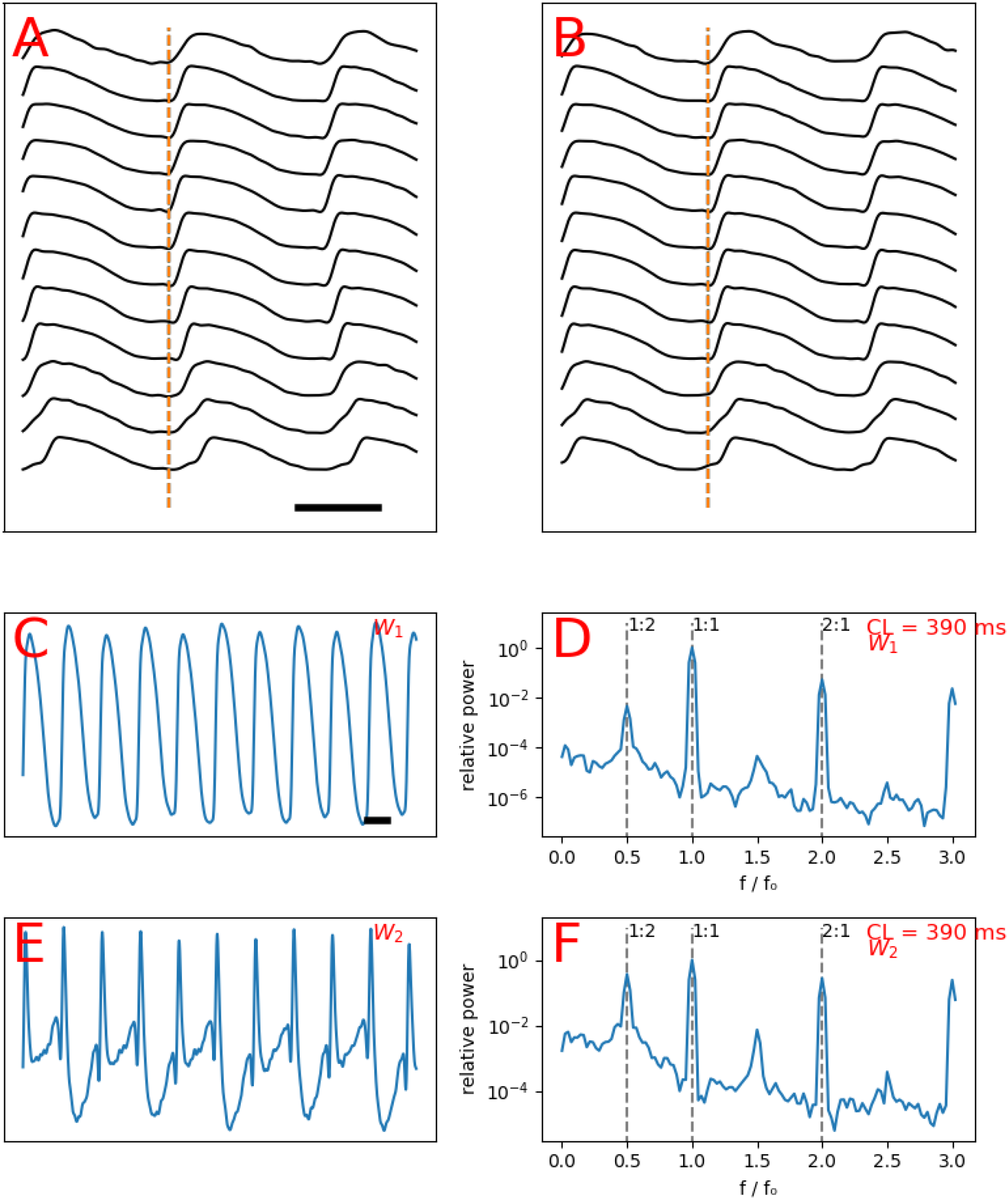
The schematics of global analysis. Spatiotemporally processed signals recorded over a line, showing staggered action potentials upstrokes consistent with wavefront propagation (**A**). Same signals as A shifted to align the upstrokes (**B**). The first principle component (*W*_1_), displaying alternans (**C**). Spectrogram of C, showing a 1:2 peak of alternans (**D**). The second principle component (*W*_2_), displaying more pronounced alternans larger than C (**E**). Spectrogram of E, showing a prominent 1:2 peak of alternans (**F**). The bar in A depicts 200 ms.

The core of our global processing routines is dimensionality reduction. We use the standard Principal Component Analysis (PCA) method. Each frame-shifted signal cube is flattened into a two-dimensional matrix (one temporal and one spatial dimension) and subjected to the truncated Singular Value Decomposition.

The top few (∼5-10) principal components capture the bulk of the dynamics (Figures 1, panels C and E show the top two principal components, marked as W1 and W2). Finally, we generate the spectrograms of the top principal components (Figures 1, panels D and F). The frequency is normalized to the frequency of the driving stimulation; therefore, the 1:1 peak corresponds to the principal action potential propagation. We are mainly interested in the sub-harmonics of the 1:1 peak. The 1:2 peak (located at exactly half the driving frequency) is a sign of period-2 alternans. Similarly, the 1:4 peak is a marker of the period-4 oscillation in the repolarization phase.

After the global stage confirms the presence of a 1:4 peak, we apply the local analysis to find higher-order periodicities (up to period-8 in this paper) for each pixel. The local analysis uses a combinatorial algorithm.^28^ The algorithm is described in Supplement B. The key idea is to assign each action potential (beat) in the input sequence to disjoint categories. For example, to detect period-2 (classic alternans), the algorithm assigns each beat to either category *A* (say, long APD beats) or category *B* (short APD beats). Then, a perfect alternating sequence is *ABABAB* …. For such a sequence, we can simply assign *A* to the odd beats and *B* to the even beats. However, the input sequence may glitch (e.g., two adjacent beats are both short APD) such that the odd/even algorithm fails to work. This problem is especially relevant to higher-order periodicity, where such glitches and frame-shifts are the rule rather than the exception. The combinatorial algorithm is designed to overcome these shortcomings. For detection of period-4, we expand the possible classes to {*A, B, C, D*}. Now, an ideal input sequence is *ABCDABCDA* …, such that all the beats assigned to *A* are similar to each other but dissimilar from other beats, and the same for beats assigned to *B, C*, and *D*. Having such an assignment, we can find the dominant periodicity of each pixel.

## Results

### The General Characteristics of the Hearts

We report on six explanted human hearts (designated **H1** to **H6**) removed during heart transplantation surgery. The general features and background of the hearts are presented in Table 1.

**Table 1:** the baseline characteristics of the hearts.

| **Heart** | **Age/Sex** | **Diagnosis** | **Amiodarone** | **Inotropes** | **LVAD** |
| --- | --- | --- | --- | --- | --- |
| **H1** | 54/M | hypertrophic cardiomyopathy | no | yes | no |
| **H2** | 38/F | non-compaction | no | yes | no |
| **H3** | 60/F | doxorubicin toxicity | yes | yes | no |
| **H4\*** | ? | ? | no | no | no |
| **H5** | 56/M | viral myocarditis | yes | yes | no |
| **H6** | 40/M | idiopathic cardiomyopathy | yes | no | no |

\* Rejected donor heart.

### Global Analysis

Three hearts (**H1, H2**, and **H3**) showed a prominent and statistically significant 1:4 peak. A borderline peak of uncertain significance was seen in another one (**H4**). No 1:4 peak was present in two hearts (**H5** and **H6**). All hearts exhibited pronounced 1:2 peaks (classic APD alternans).

The two hearts (**H1, H2**) with the most prominent 1:4 peak were not on a membrane-active antiarrhythmic medication, whereas the two without a 1:4 peak (**H5, H6**) were on amiodarone (Table 1).

Figure 2 (panel A) depicts the global spectrogram of **H1**. The baseline spectrogram at 500 ms shows the expected 1:1 peak (the primary activation) and a small 1:2 peak, signifying repolarization alternans. As the heart was stimulated faster at a cycle length of 310 ms, the 1:2 peak became larger (higher amplitude alternans), and a new 1:4 peak emerged. This means that a bifurcation occurred somewhere between 500 ms and 310 ms, and the dynamics had period-4 periodicity. Pacing this heart faster at 300 ms resulted in VF.

**2.**
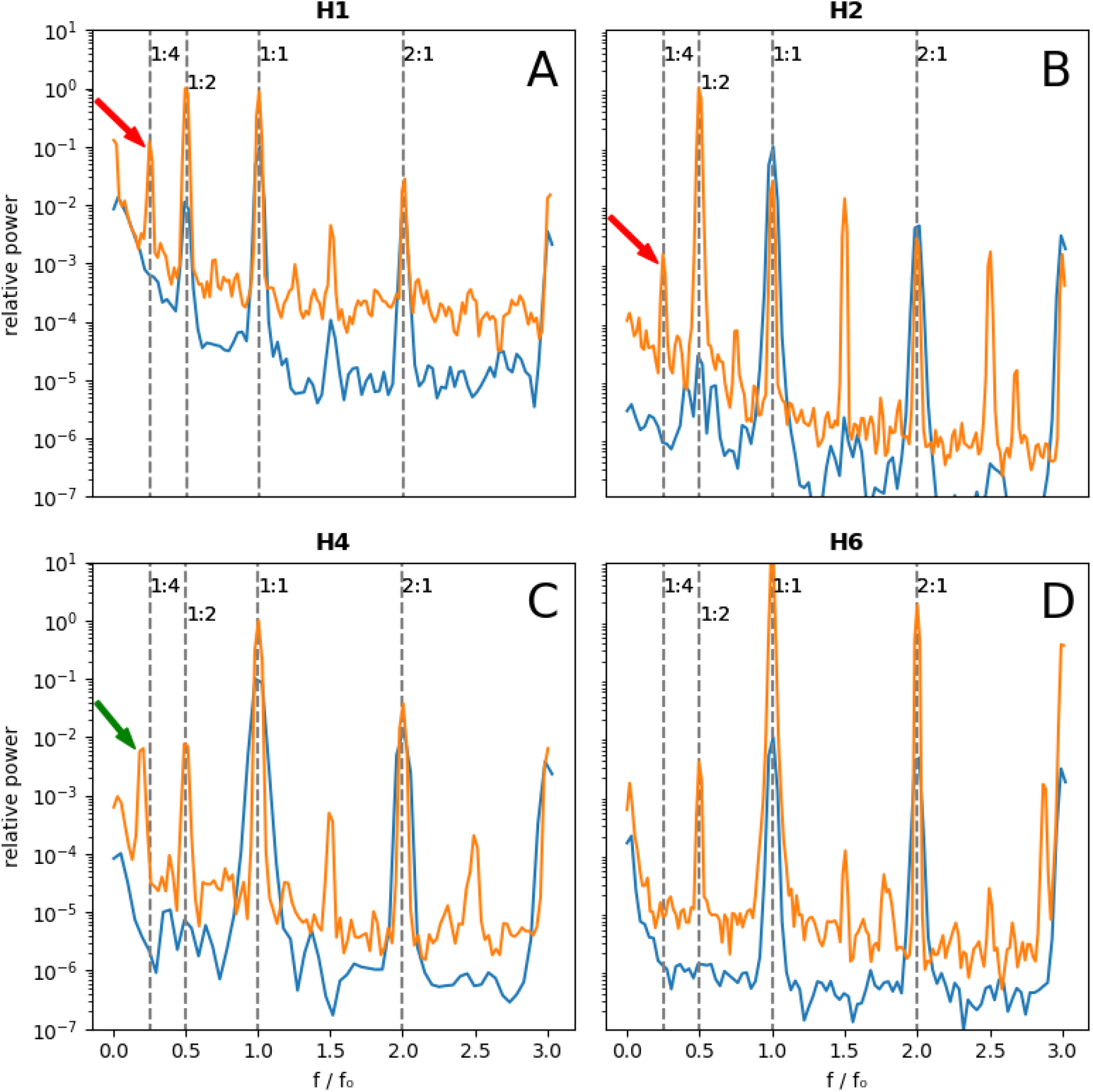
Comparison of baseline and pre-VF spectrograms using global analysis. The blue spectrograms are the baseline (stimulation cycle length of 500 ms except for **H4** at 800 ms) and the orange spectrograms are obtained just prior to VF induction. **H1, H2** (**A and B**) exhibit prominent 1:4 peaks (the red arrows), while no discernable 1:4 peak is seen for H6 (**D**). **H4** has a ∼0.18 peak (the green arrow), corresponding to mainly period-6 activity **(C)**, which is discussed in the text. The baseline signals are multiplied by 0.1 to offset the signals for better visualization.

**H2** follows a similar pattern (panel B). Again, we observed barely discernible alternans at 500 ms with a strong 1:2 peak and a clear 1:4 peak at 270 ms. Similarly, VF was induced while pacing at 260 ms.

The peak of interest in **H5** (panel C) is offset from the 1:4 location and is around ∼0.18. This peak is further discussed below.

**H6** is a negative example with no 1:4 peak (panel D). There is a prominent 1:2 alternans peak while stimulating at 270 ms, but no significant peak at 1:4. Stimulating this heart faster resulted in a conduction block but no reentrant arrhythmia. As mentioned above, this heart was on amiodarone. Interestingly, when the experiment was repeated at room temperature instead of the usual 37 C, a weak 1:4 peak appeared (data not shown).

The absence of the 1:4 peak at 500 ms in hearts with a prominent 1:4 peak at short cycle lengths significantly reduces the chance that this peak is a processing artifact and points to its dynamic origin. We can probe the dynamics further by looking at the stimulation frequency dependencies of the 1:2 and 1:4 peaks (see Supplement C).

### Local Analysis

The distribution of areas with higher-order periodicity is heterogeneous in both space and time. Figure 3 panel A shows the results of local analysis applied to **H1** superimposed on the activation (isochronous) map. Stable and mostly static period-4 periodicity is localized to a few discrete islands with roughly circular borders. These islands correspond to anatomical structures like papillary muscles. However, higher-order dynamics is not limited to these islands. Movie S1 (see Supplement D) shows the dynamics of transient higher-order (up to period-8) periodicities. This movie visualizes the spontaneous period for each pixel and at each beat. A typical frame from the movie is reproduced in panel B. Furthermore, sample signals (panels E and G) and the APD trends (panels F and H) from the stable 1:4 areas are presented and compared to a control point (panels C and D) with only 2:1 alternans.

**3.**
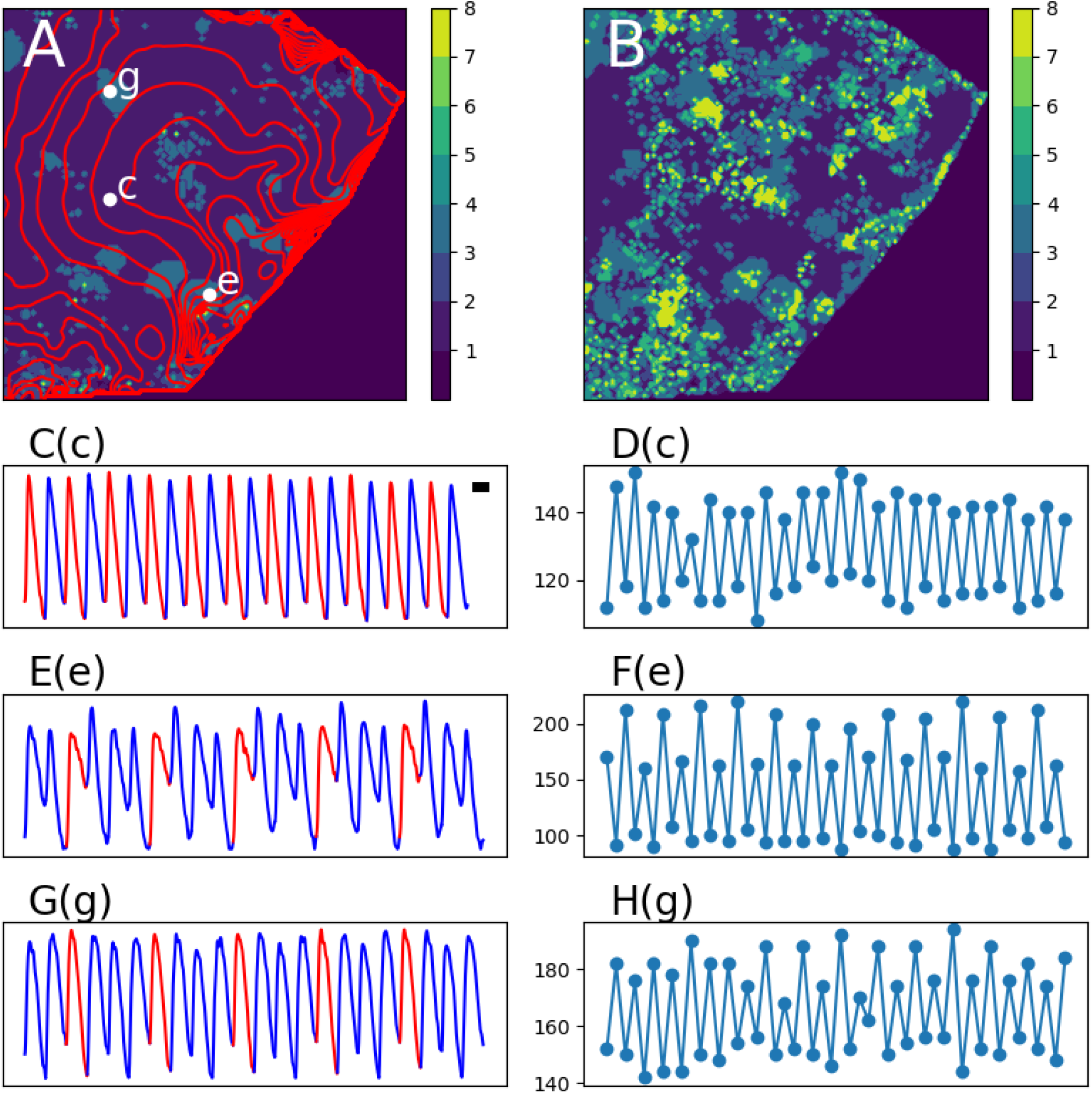
Local analysis of heart H1. The predominant periodicity of the pixels (color coded) superimposed on an activation map (red isochrones)(**A**). A frame from Movie S1, showing the instantaneous period of each pixel (**B**). The representative signals from points **c** (period-2), and **e** and **g** (period-4)(**C, E, G**) with the corresponding APD trends (**D, F, H**). Note that stable period-4 is localized to a few discrete regions. The isochronous lines are 10 ms apart and the bar in C is 100 ms long.

Figure 4, depicting **H2**, is similar. Stable period-4 islands are close to slow zones, as deduced from isochrones crowding, and anatomical structures (panel A). In addition, there are two 1:6 areas. The spatiotemporal higher-order dynamics is shown in Movie S2 (a representative frame in panel B). Higher-order dynamics occurs over arcs, which are parallel to the isochrones. The APD trend from a representative 1:6 area (panel H) is aperiodic, in contrast to the 1:2 and 1:4 APD trends.

**4.**
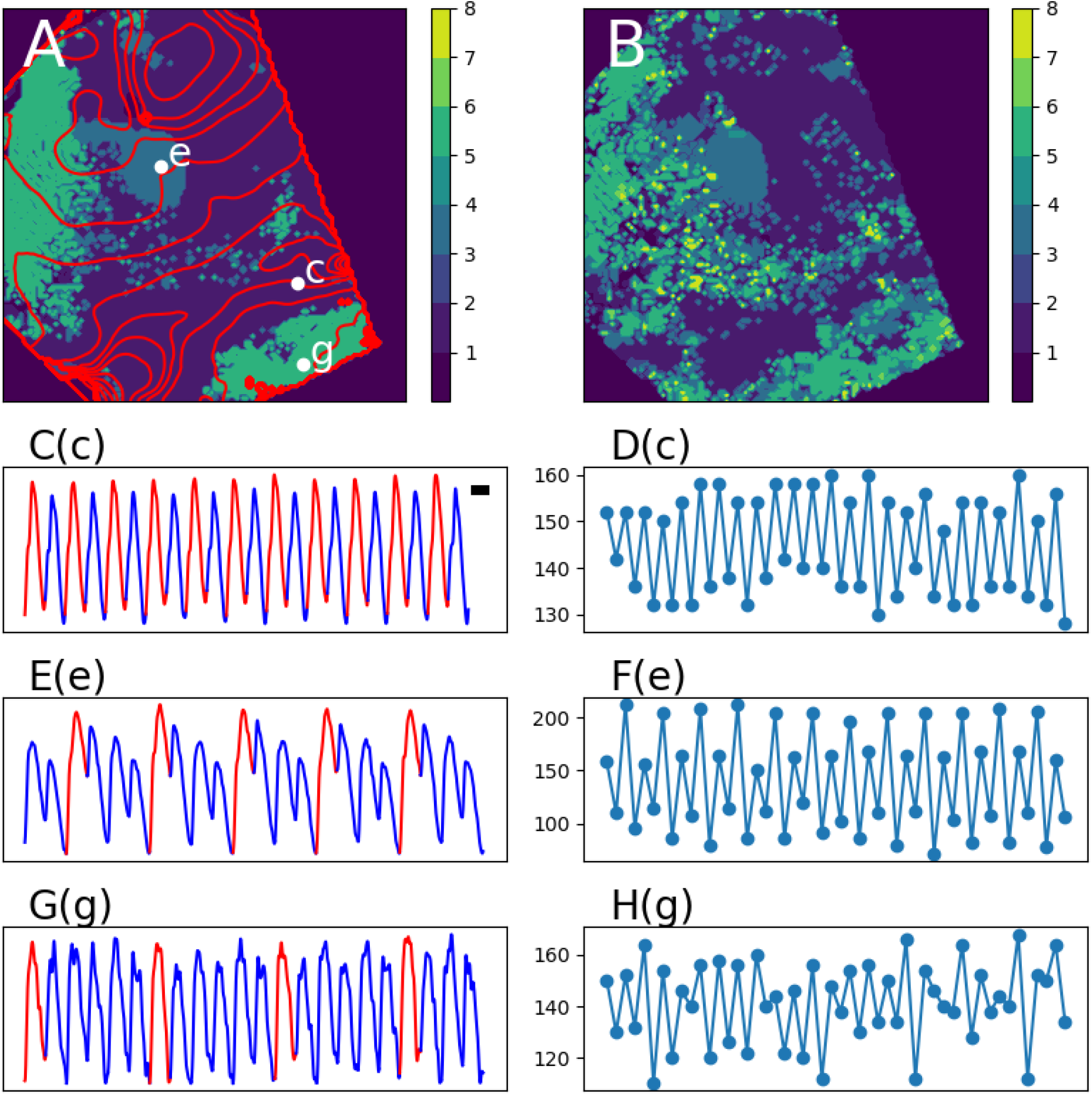
Local analysis of heart H2. The predominant periodicity of the pixels (color coded) superimposed on an activation map (red isochrones)(**A**). A frame from Movie S2, showing the instantaneous period of each pixel (**B**). The representative signals from points **c** (period-2), **e** (period-4) and **g** (period-6)(**C, E, G**) with the corresponding APD trends (**D, F, H**). Period-6 areas seem disjoint from period-4 regions. The isochronous lines are 10 ms apart and the bar in C is 100 ms long.

Figure 5 is from **H5** with no 1:4 peak but a prominent 0.18 peak (Figure 2, panel C). The local analysis shows a baseline of 1:2 periodicity mixed with expanses of predominantly 1:6 periodicity and a small 1:5 area. Movie S3 and the corresponding frame in panel B confirm that the dominant dynamics is 1:6. Like **H2**, the APD trends from the 1:5 and 1:6 areas are aperiodic (panels F and H). Similarly, the higher-order areas, especially the 1:8 regions, form arcs parallel to the isochrones.

**5.**
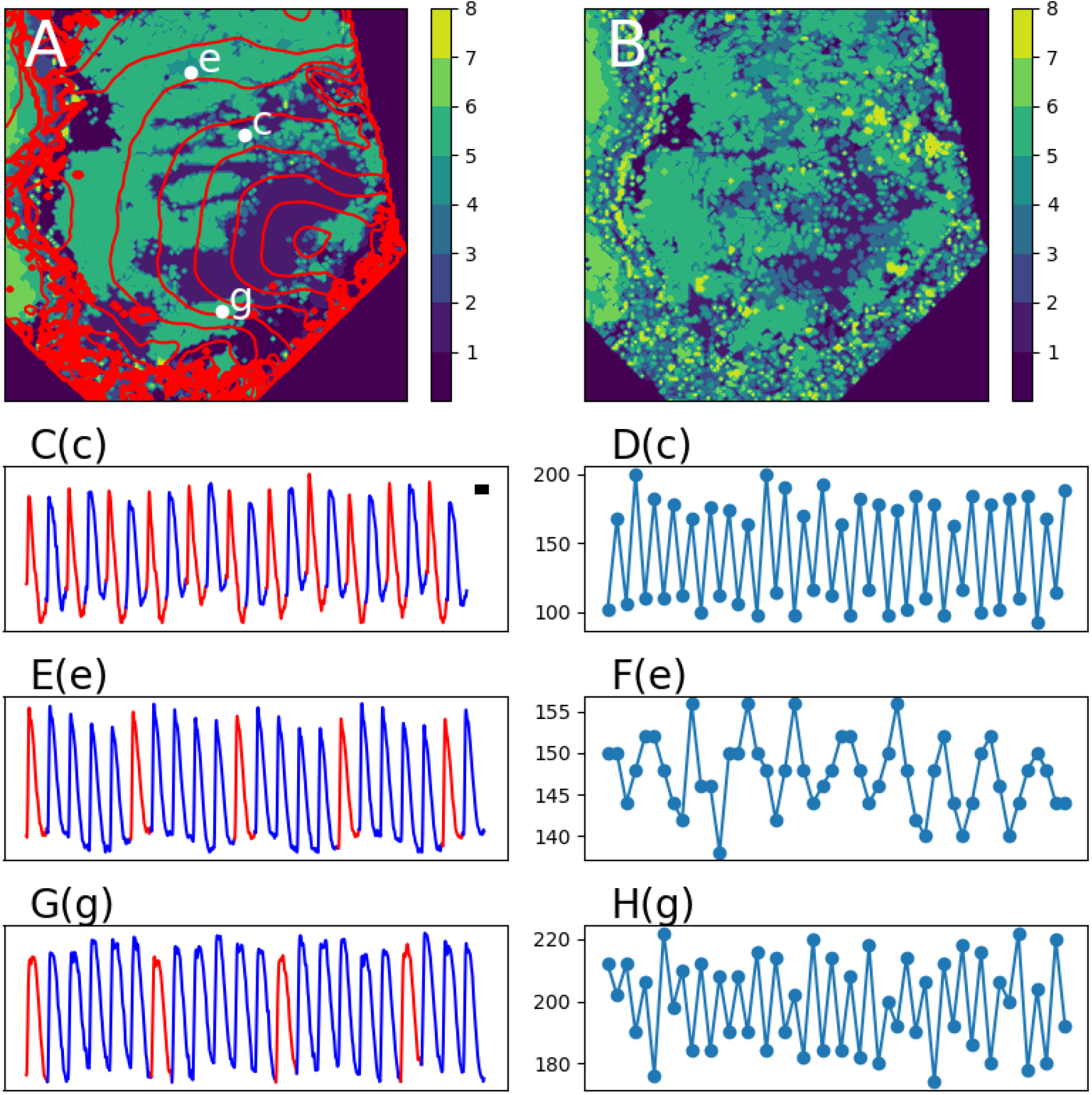
Local analysis of heart H4. The predominant periodicity of the pixels (color coded) superimposed on an activation map (red isochrones)(**A**). A frame from Movie S3, showing the instantaneous period of each pixel (**B**). The representative signals from points **c** (period-2), and **e** (period-5) and **g** (period-6)(**C, E, G**) with the corresponding APD trends (**D, F, H**). No significant regions with period-4 are seen in this heart that lacks a 1:4 peak in global analysis. The isochronous lines are 10 ms apart and the bar in C is 100 ms long.

## Discussion

Repolarization alternans is one of the bedrocks of theoretical cardiac electrophysiology. It manifests at the cellular level as the APD alternans and at the tissue level as TWA. Alternans importance stems from the link that it provides between cellular dynamics and fibrillation at the organ scale. This paper looks beyond period-2 alternans to higher-order dynamics in human hearts.

We report what we believe to be the first detection of stable period-4 and intermittent higher order periodicities in human cardiac tissue during fast stimulation. Periods of order 2*n* are stable and consistent with the expected behavior of period-doubling. On the other hand, we also observed areas of transient period-5 and period-6. We interpret period-6 as a combination of APD alternans with an underlying period-3 oscillation. Famously, period-3 implies chaos according to Sharkivsky’s theorem.^29^ The detection of period-5 and period-6 with the corresponding aperiodic APD trends strongly suggests the existence of pockets (tissue regions) of chaos during fast pacing but before VF.

These findings have implications for the mechanisms of VF initiation. According to the concordant to discordant alternans pathway, the heart is in a meta-stable but not chaotic condition just before VF initiation, waiting for a trigger to tip the balance. The trigger is usually a conduction block or a premature ventricular contraction caused by either early after depolarization (EAD) or phase-2 reentry.^30,31^ On the other hand, based on the period-doubling mechanism, as the heart is stimulated at faster rates, higher periods occur heterogeneously until some areas become chaotic. These chaotic regions may spatially grow to act as niduses of instability that then degenerate regular rhythms into chaotic fibrillation. The distinction between the two mechanisms is, first, the coexistence of regional chaos with non-chaotic areas prior to VF (observed in this study), and second, a causal link between the chaotic regions and VF initiation (not enough data).

It should be noted that the two mechanisms of VF initiation, through discordant alternans or period-doubling, are not mutually exclusive. For example, the chaotic regions may produce the premature beat that triggers the transition from discordant alternans to VF.

We observed that the higher-order regions were spatially heterogeneous and preferred certain areas of the heart. However, their orientation was roughly parallel to the isochrone lines. These observations suggest that higher-order dynamics are generated by a mixture of static (related to the pre-existing cellular electrophysiological properties) and dynamics (patterns caused by the interplay of the repolarization and depolarization dynamics) mechanisms.

Our data points to the different effects of membrane-active antiarrhythmic medications on period-4. In the absence of exposure to antiarrhythmic medications, the 1:4 peak was taller and more prominent. Conversely, we did not observe the 1:4 peak in the hearts treated with amiodarone. While our small sample size precludes a statistical statement, this result aligns with the dynamical predictions. Specifically, the disjoint effect of amiodarone on periods of order 2*n* compared to period-6 suggests different dynamical mechanisms and warrants further study.

The detection of higher-order periodicities in human hearts may have practical implications in addition to providing mechanistic insights. If a practical method to detect period-4 from clinical recordings (e.g., surface ECG) can be developed, it may fix the main shortcoming of T wave/APD alternans in the form of low positive predictive value for malignant ventricular arrhythmias.^11^ However, it should be noted that this is potentially a very difficult task, considering the extent of signal processing required to detect higher-order oscillations and the rarity of prior experimental evidence of high-order oscillations in cardiac tissue.

Ablation, i.e., delivery of radiofrequency or other energy sources to damage and kill cardiac cells in strategic locations with the goal of disrupting arrhythmia circuits, is one of the core therapies for sustained VT but has a minimal role in VF due to the dynamical nature of VF. If it is confirmed that higher-order periodicities have a causal link with VF induction and there is a significant static component to their localization, it is possible that ablation of these static areas can help to prevent VF.

Hodgkin-Huxley-type ionic models are commonly used to simulate cardiac electrophysiology. Hundreds of such modes are proposed to explain various cell types in different species.^32^ However, many of these models fail to reproduce the correct short cycle length behavior and have abnormal APD alternans characteristics. The emergence of higher-order dynamics and their potential suppression by membrane-active antiarrhythmic medications can serve as a stringent test to select physiologically accurate models for the study of arrhythmias and their prevention, control and termination.

## Acknowledgements

The study protocol was approved by the Emory University and Georgia Institute of Technology Institutional Review Boards (IRB) under Protocol Number H22204.

## Sources of Funding

This study was supported in part by the NIH under grant 1R01HL143450-01 and the NSF under CMMI-1762553.

## Disclosures

Authors have no disclosure to make.

## Supplement Materials

**Supplement A:** Spatial Filtering

**Supplement B:** Local Analysis

**Supplement C:** The stimulation frequency dependencies of the 1:2 and 1:4 peaks

**Supplement D:** Movies of beat-to-beat periodicity

**Movie S1:** beat-to-beat periodicity of **H1**

**Movie S2:** beat-to-beat periodicity of **H2**

**Movie S3:** beat-to-beat periodicity of **H5**

## Notes

### Competing Interest Statement

The authors have declared no competing interest.

